# Anion efflux mediates transduction in hair cells of zebrafish lateral line

**DOI:** 10.1101/2022.07.11.499370

**Authors:** Elias T. Lunsford, Yuriy V. Bobkov, Brandon C. Ray, James C. Liao, James A. Strother

## Abstract

Hair cells are the principal sensory receptors of the vertebrate auditory system, and transduce sounds with mechanically-gated ion channels that permit cations to flow from the surrounding endolymph into the cells. The lateral line of zebrafish has served as a key model system for understanding hair cell physiology and development, and it has often been speculated that these hair cells employ a similar transduction mechanism. In this study, we demonstrate that the hair cells are exposed to an unregulated external environment with cation concentrations that are too low to support transduction. Instead, our results indicate that hair cell excitation is mediated by a fundamentally different mechanism involving the outward flow of anions.

## Main

Vertebrate hair cells are ciliated mechanoreceptive sensory cells responsible for the exquisite sensitivity of the auditory and vestibular systems. In the auditory system, these cells are immersed in endolymph with a relatively high K^+^ concentration and positive potential, which establishes a strong electrochemical gradient favoring K^+^ influx into the cells^1–3^. Sound propagating through the cochlea generates shearing forces on the apical stereocilia bundle of hair cells. Deflection of the stereocilia opens mechanically-coupled cation channels (i.e., TMC1/2^4–6^), which permits K^+^ influx into the hair cells (mechanoelectrical transduction [MET] current) and drives membrane depolarization^7,8^. The high K^+^ concentration of the endolymph is key to this pathway, and is produced by active secretion of K^+^ into the endolymph by strial marginal cells via a process that includes K^+^ channels and Na^+^/K^+^ ATPase^9–11^.

The hair cells of the inner ear are thought to have evolved from anatomically similar cells found in the lateral line of fishes and amphibians (reviewed by^12^). The lateral line system detects the movement of water around the body and is critical for survival, as it mediates behaviors such as predator avoidance, prey capture, and navigation^13–16^. Unlike the hair cells of the inner ear, the lateral line hair cells lie on the surface of the body and are surrounded by the external environment rather than a K^+^-rich endolymph. Work in amphibians in the 1970s indicated that the gelatinous cupula that encapsulates the lateral line hair cells maintains an ionic microenvironment comparable to the inner ear endolymph, which establishes the ionic gradient necessary for cation influx mediated mechanotransduction^17,18^.

Although freshwater zebrafish are a powerful model system for understanding hair cell physiology^19–21^, this foundational hypothesis has remained largely untested. Here, we show that the ionic microenvironment in the cupula of the superficial neuromasts of zebrafish larvae is indistinguishable from the surrounding freshwater. Electrochemical calculations indicate that the freshwater inhabited by zebrafish does not provide ionic gradients sufficient to support the putative cation-mediated mechanotransduction mechanism. Instead, our results suggest a novel process driven by anion efflux. For cells in contact with ion-poor extracellular saline, typical negative resting membrane potentials and intracellular Cl^-^ concentrations are sufficient to generate anion efflux capable of inducing robust membrane depolarization. Although studies of sensory physiology often focus on cation mediated transduction process, anion efflux has been observed to contribute to signal amplification in vertebrate chemosensory receptors, which often directly interface with an unregulated external environment^22–26^. It has also been argued that anion efflux may confer favorable properties, including reduced sensitivity to large variations in extracellular ionic composition^27,28^.

### The cupula does not provide a cation-rich microenvironment for lateral line hair cells

We first examined if the cupula of the superficial neuromasts contained an elevated K^+^ concentration capable of supporting mechanotransduction. Animals were bathed in a freshwater media (E3 saline) with a K^+^ selective fluorescent indicator (IPG-4). Using confocal microscopy, we found that fluorescence in the cupula was nearly indistinguishable from that of the surrounding water, suggesting that the cupula does not maintain elevated K^+^ concentrations (Fig. 1B). To quantify the K^+^ concentration within the cupula, the indicator was saturated by adding K^+^ to the surrounding media, and the recorded fluorescence values were fit to a model that incorporates both the binding affinity of the indicator and potential differences in indicator concentration between the cupula and media. Consistent with our qualitative observations, the calculated K^+^ concentration of the cupula was not significantly different from the surrounding media (p = 0.49, N=14; Fig. 1C). Analogous experiments were then performed examining each of the major cations present in freshwater saline and again no significant differences were found between the cupula and the surrounding water (Na^+^: p = 0.23, N=9; Ca^2+^: p = 0.54, N = 9; H^+^: p = 0.38, N=9; Fig. 1D). We also examined the Cl^-^ concentration within the cupula, since it is the likely counterion for any cation, and found that it was also not significantly different from the freshwater media (p=0.43, N=6, Fig. 1D). These results indicate that the cupula does not support an ionic microenvironment, and the apical surfaces of the lateral line hair cells are directly exposed to an ion-poor freshwater saline that is markedly different from the endolymph of the inner ear^2,3,29–31^.

**Fig. 1.**
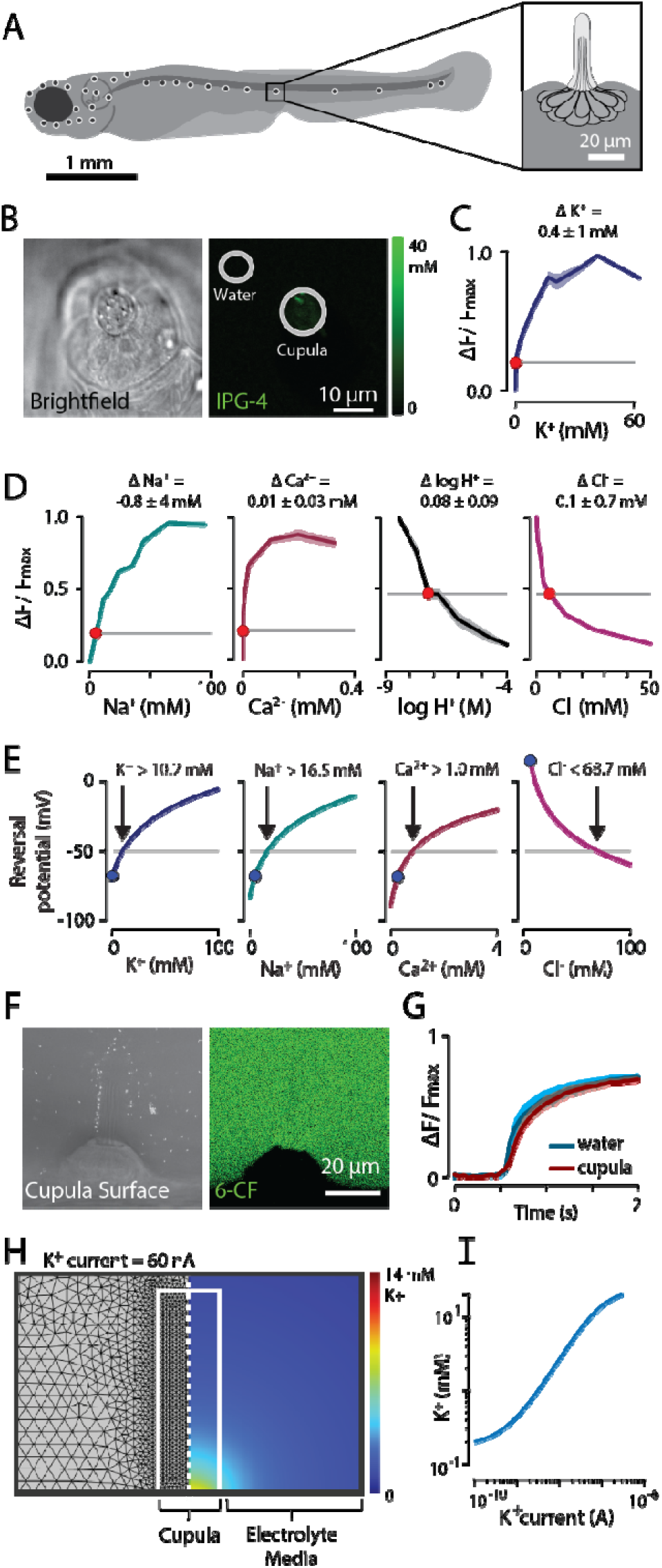
The ionic microenvironment of the cupula resembles freshwater and cannot support cation influx mediated depolarization. **(A)** Superficial neuromasts of the lateral line system in larval zebrafish. **(B)** *In vivo* visualization of K^+^ in the vicinity of a neuromast using the fluorescent probe IPG-4 (left: brightfield; right: IPG-4). **(C)** Fluorescence of K^+^ probe in solutions with varying K^+^ concentrations (blue curve), fluorescence expected for an environment matching surrounding freshwater media (E3 saline; horizontal grey line), and measured fluorescence within the cupula (red dot, error bars are ± SEM). Using these values, the difference between the K^+^ concentration in the cupula and the surrounding media was calculated (ΔK^+^, mean ± SEM). **(D)** Results from experiments similar to (C) except examining Na^+^ (probe: ING-2), Ca^2+^ (probe: Fluo-5N), H^+^ (probe: BCECF), and Cl^-^ (probe: MQAE). In all cases, ion concentrations within the cupula matched those of the surrounding media. **(E)** Predicted reversal potentials for MET channels as a function of cupula cation concentration. The estimated opening potential of voltage-gated Ca^2+^ channels at the basal membrane is indicated (−50 mV, horizontal line). The cation concentration necessary to reach this potential (black arrow) and the concentration in freshwater media (blue dot) are also indicated. The predicted reversal potential for a hypothetical chloride channel is also shown (right). **(F)** Images showing penetration of a charged fluorophore (6-carboxyfluorescein, 6-CF) into neuromast cupula (left: cupula surface visualized with microspheres, right: 6-CF). **(G)** From the same experiments as (F), time series showing rapid increase in fluorescence in both media around the cupula and within the cupula after introduction of the charged fluorophore. **(H)** Simulation of a hypothetical K^+^ microenvironment, produced by solving 3D Nernst-Plank equations (left: mesh, right: K^+^ distribution for K^+^ current = 60nA). **(I)** K^+^ concentration at apical surface of hair cells as a function of K^+^ secretion current.

To systematically explore the consequences of the hair cells being exposed to an ion-poor environment, we calculated the membrane reversal potential for a MET channel in different extracellular solutions, assuming relative permeabilities similar to other hair cells. We found that in freshwater media (E3 saline), MET conductance would be expected to produce K^+^ efflux and hyperpolarization rather than depolarization, which strongly suggests that K^+^ current is not the primary driver of transduction (Fig. 1E). MET conductance is only predicted to induce membrane depolarization sufficient to open the voltage-gated Ca^2+^ channels on the basolateral membrane (Ca_V_1.3^32^) in water with cation concentrations much greater than those in the natural habitat of zebrafish (Fig. 1E^29–31^). Although the chemical gradient favors Ca^2+^ influx into the hair cell, this influx is unable to overcome the hyperpolarizing effect generated by K^+^ efflux given the estimated relative permeabilities of the MET channels. Ca^2+^ currents would therefore only be sufficiently depolarizing if the MET channels displayed Ca^2+^ selectivity far in excess of that reported in other systems^4^. We next calculated the reversal potential for a hypothetical Cl^-^ channel and found that Cl^-^ conductance would be expected to readily drive Cl^-^ efflux from the hair cells and membrane depolarization in a wide range of external environments (Fig. 1E). In total, these results argue against K^+^ influx as a mechanism for lateral line hair cell depolarization and suggest that Ca^2+^ influx or Cl^-^ efflux may have more central roles.

The lack of an ionic microenvironment in the cupula would profoundly affect the mechanotransduction mechanisms available to hair cells. We next conducted a series of experiments and simulations further exploring whether the physical properties of the cupula could support such a microenvironment. We first examined the diffusion of charged molecules within the cupula by rapidly introducing a negatively charged small molecule fluorophore (6-carboxyfluorescein; 6-CF) around the cupula and imaging its diffusion from the media into the cupula (Fig. 1F). There were no significant differences between the rate at which fluorescence increased in the cupula and the surrounding water, suggesting that charged molecules rapidly penetrate the cupula (relative time constant = 0.88 ± 0.06 [mean ± S.E.M.], p = 0.076, N=9, Fig. 1G). Computational simulations confirmed this experimental setup had sufficient resolution to detect meaningful differences in the diffusive properties of the cupular matrix (Extended Data Fig. 1). This finding is also consistent with empirical studies of the ultrastructure of lateral line cupulae, which identified no membrane or other dense surface structure^33,34^.

We next evaluated the ion currents that would be necessary to sustain a K^+^ reservoir in the cupula using computational simulations. Since this configuration violates the assumptions of a typical neuron electrical equivalent circuit^35^, we simulated these dynamics by directly solving the full three-dimensional Nernst-Planck equations using the finite element method (Fig. 1H). Based on the above diffusion experiments, we estimated the diffusion coefficients for ions moving through the cupular matrix to be similar to water and assumed a constant rate of K^+^ secretion into the matrix at the base of the neuromast. We found that a K^+^ current 56 nA would be needed to produce the K^+^ concentration of 10.2 mM required for depolarizing MET conductance (Fig. 1E&I). Although the ionoregulatory currents of the neuromast have not been quantified, these K^+^ current values are several orders of magnitude greater than the resting MET current of hair cells^36,37^. These results indicate that recently described skin-derived ionocytes are unlikely to maintain a K^+^ microenvironment at the apical surface of the hair cells, but may instead support regulation of the extracellular environment at the basolateral surfaces^38^.

Cumulatively, these experiments indicate that the cupula does not maintain a cation rich microenvironment sufficient to support hair cell depolarization, challenging a long-standing assumption that hair cell transduction is cation driven in the lateral line system of zebrafish and other freshwater fishes.

### Lateral line function only requires micromolar extracellular calcium

What other mechanisms could mediate hair cell depolarization? We next tested the hypothesis that Ca^2+^ could contribute to hair cell depolarization, given its reversal potential and assuming MET channels with unusually high Ca^2+^ selectivity. Hair cells expressing the fluorescent calcium indicator GCaMP6s were imaged while mechanically deflecting the cupula with a piezoelectric transducer (Fig. 2A). Changes in fluorescence near the basal surface of the hair cells were observed, which have been attributed to Ca^2+^ influx through voltage-gated channels on the basal membrane^21^. Hair cells produced robust responses to this stimulus in freshwater media (E3 saline, 330 μM Ca^2+^; p < 0.001, df = 27; Fig. 2A). Interestingly, we found that hair cells continue to respond in media with substantially reduced Ca^2+^ concentrations (20 μM Ca^2+^; p = 0.04, df = 3; Fig 2A) and that these responses were indistinguishable from those observed in freshwater media (p = 0.13, df = 3). The insensitivity of hair cell responses to available Ca^2+^ suggests that Ca^2+^ is not the principal carrier of the MET current. However, responses in 0 μM Ca^2+^ media were significantly reduced compared to those in freshwater media (p = 0.02, df = 9; Fig 2A). This indicates that environmental Ca^2+^ modulates MET currents in naturalistic freshwater media, consistent with observations of Ca^2+^ effects on hair cell mechanotransduction in zebrafish and other species^8,39–43^.

**Fig. 2.**
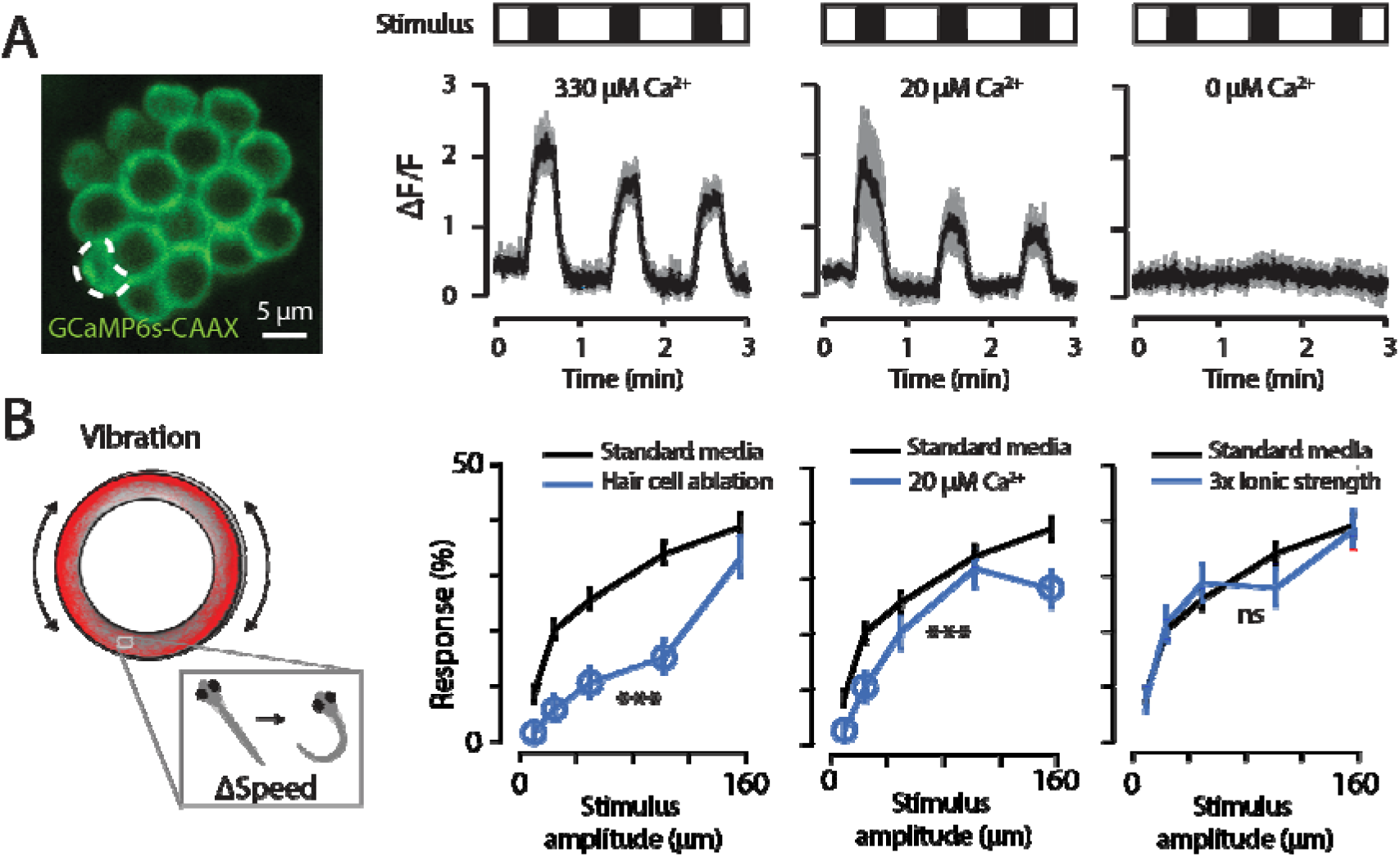
Only micromolar extracellular calcium is required for lateral line function. **(A)** Response of hair cells to a mechanical stimulus, recorded by imaging genetically expressed fluorescent calcium indicator GCaMP6s-CAAX at the basal membrane (left: typical image, region of interest indicated; right: responses in 330 μM, 20 μM, and 0 μM Ca^2+^, mean ± SEM). Hair cell responses in micromolar Ca^2+^ media are comparable to those in freshwater media (E3 saline), while responses are abolished in calcium-free media. **(B)** Behavioral response of zebrafish larvae to a mechanical stimulus under varying conditions, recorded with a novel behavioral assay that employs an oscillatory Couette cell to produce a pure-shear flow stimulus (left: schematic of behavioral setup with typical trajectory in red; left: probability of response following neomycin-induced hair cell ablation, in 20 μM Ca^2+^ media, and in 3x ionic strength media). Asterisks indicate statistically significant differences integrated across stimulus intensities (*p<0.05, **p<0.01, ***p<0.001), open circles represent significant pairwise comparisons (p<0.05), and results are shown as mean ± SEM.

To verify that these cellular-scale differences are reproduced at the organismal level, we next examined the effects of low Ca^2+^ environments on the behavior of freely swimming larvae. A number of behavioral assays have been successfully employed for assessing lateral line function in zebrafish larvae, including assays measuring C-start escapes in response to impulsive stimuli and assays measuring rheotactic responses^13–15^. However, the contribution of multiple sensory modalities (e.g. vestibular, acoustic, visual) to behavioral responses often presents a challenge when developing such assays. Since the superficial lateral line is principally sensitive to shear at the body surface^44^, we designed a novel behavioral assay that uses an oscillatory Couette cell to produce a shearing flow without accompanying pressure waves (Extended Data Fig. 2). The position and swimming speed of animals were continuously monitored, and stimuli of varying intensities were presented at random intervals. Each stimulus that produced a significant change in swimming speed (outside 95% confidence interval) was recorded and then the net response probability was calculated as a function of stimulus intensity. In order to reduce both habituation and activation of collateral sensory modalities, we intentionally examined small stimulus intensities that produced modest changes in behavior with relatively low response probabilities rather than escape responses with high probability.

To verify the efficacy of this assay, we confirmed that animals responded in an intensity-dependent manner and that neomycin-induced hair cell ablation dramatically decreased this response (Fig. 2B). Next, we examined how low environmental Ca^2+^ affects lateral line sensitivity and found that animals displayed reduced but robust responses even with Ca^2+^ decreased by >15x (21% decrease with 50 μm stimulus, omnibus p < 0.001, N = 30 exp/94 ctrl). These results are consistent with our imaging data and data from other species^39,45^. Using this same assay, we then examined how lateral line sensitivity is altered by increases in the ionic strength of the freshwater environment. The responses of animals in media with a 3x increase in ionic strength were not significantly different from those in freshwater media (E3 saline, omnibus p > 0.05, N=30 exp/94 ctrl). Media with higher ion concentrations were also explored, and significant decreases in sensitivity were detected, but such ion concentrations far exceed the values observed in the natural habitat of zebrafish and these effects may result from osmotic stress rather than a specific interaction with the lateral line (Extended Data Fig. 3). In total, these findings strongly suggest that extracellular Ca^2+^ modulates the mechanotransduction process but is not the primary ion current driving hair cell depolarization.

### Calcium-activated anion efflux contributes to hair cell depolarization

Since the electrochemical gradient between hair cells and freshwater strongly favors anion efflux, we next examined whether this efflux could be contributing to mechanotransduction. We began by leveraging available transcriptome data to generate testable hypotheses. We analyzed published transcriptome scRNA-seq data collected from zebrafish lateral line neuromasts ^46^ for cells that expressed a hair cell marker (tmc2b), filtered transcripts for genes associated with ion channel activity, and then reviewed the resulting list for genes with anion efflux activity. This analysis indicated that the lateral line hair cells may express the calcium-activated anion channel Anoctamin-2b (ANO2b), also known as TMEM16b (Fig. 3A).

**Fig. 3.**
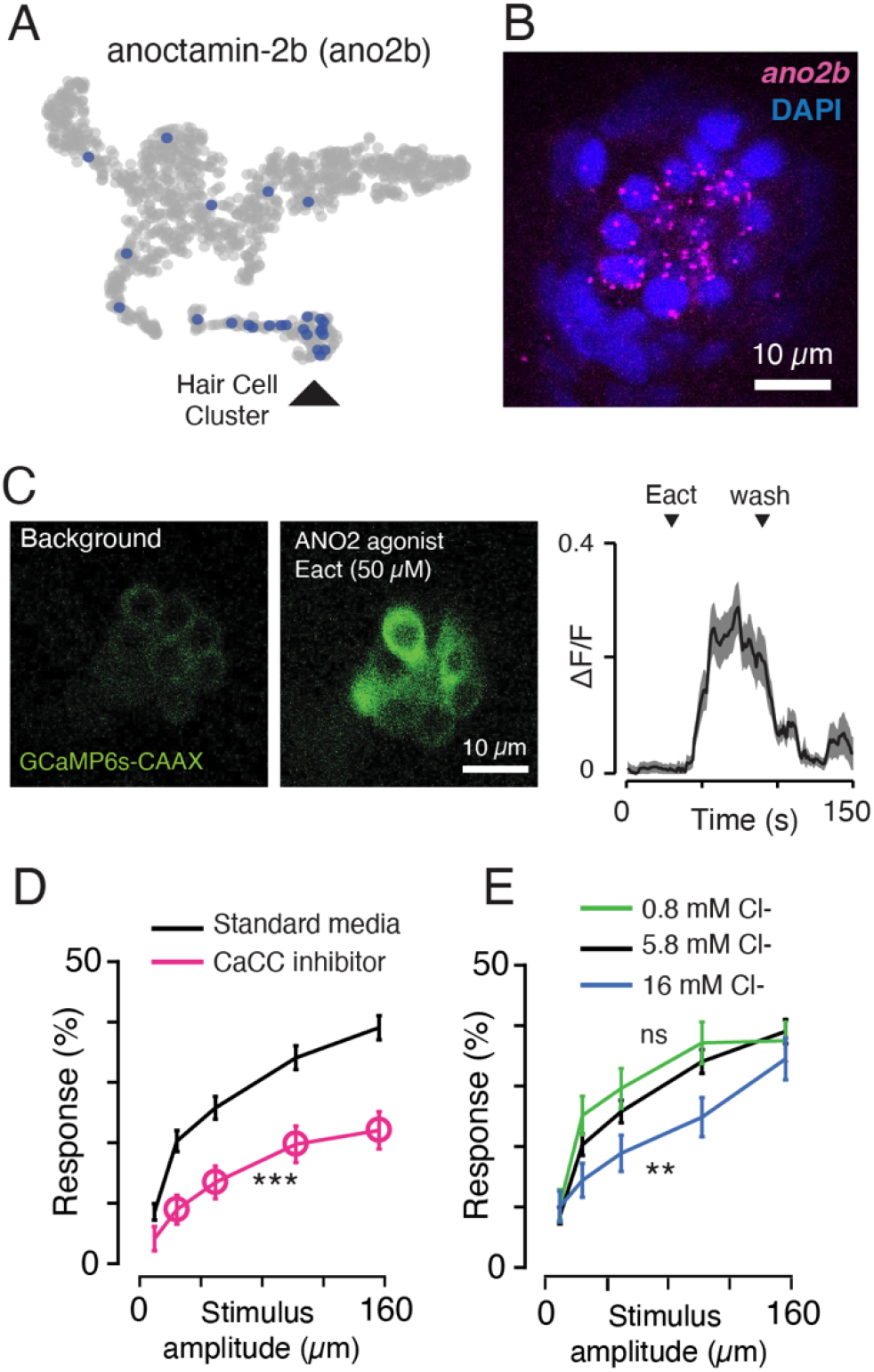
Calcium-activated chloride channels are expressed in lateral line hair cells, sufficient to activate hair cells, and necessary for functional responses at the organismal level. **A**) Visualization of single cell transcriptome data for lateral line neuromast showing expression of anoctamin-2b (*ano2b*) in the hair cell cluster (t-SNE plot, data from^46^). (**B**) *In situ* hybridization image showing *ano2b* expression in a lateral line neuromast (punctate pattern consistent with single-molecule RNA-FISH). (**C**) Response of hair cells to the Ano2 agonist Eact, recorded at the basal membrane using a genetically expressed fluorescent calcium indicator (left and middle: typical images, right: measured ΔF/F for introduction and washout, mean ± SEM.). (**D**) Effect of the calcium-activated chloride channel (CaCC) inhibitor NFA on the behavioral response of zebrafish larvae to a mechanical stimulus. Asterisks indicate statistically significant differences integrated across stimulus intensities (*p<0.05, **p<0.01, ***p<0.001), open circles represent significant pairwise comparisons (p<0.05), and results are shown as mean ± SEM. **(E)** Similar to (D), except showing the effect of decreasing the electrochemical gradient supporting chloride efflux by increasing extracellular chloride.

To verify this expression, *ano2b* transcripts were labeled using HCR RNA-FISH. We observed *ano2b* labeling in lateral line hair cells, olfactory epithelium, and the dorsal habenula (Fig. 3B & S4). The expression of anoctamin in the olfactory epithelium is consistent with similar findings in mammalian olfactory epithelium^47^, and *ano2b* expression in the dorsal habenula of zebrafish has been previously reported^48^.

We next examined if calcium-activated chloride channels are capable of inducing depolarization of the lateral line hair cells. Using zebrafish larvae that express a fluorescent calcium indicator in hair cells (*myo6b:GCaMP6s-CAAX*^21^), we recorded hair cell responses to the Anoctamin2 agonist Eact^49^. This agonist induced immediate and robust increases in intracellular calcium at the basolateral membrane (p < 0.001, df = 58), which were reversible upon washout (Fig. 3C). These responses are consistent with the hypothesis that calcium-activated anion channels contribute to mechanotransduction in the lateral line hair cells.

We then sought to determine if these effects were reflected in the responses of an intact, behaving animal. When we examined the behavioral response of animals that had been treated with the calcium-activated chloride channel blocker niflumic acid (NFA)^50^, we found that they exhibited a marked decrease in behavioral responses relative to untreated animals (48% decrease with 50 μm stimulus, omnibus p < 0.001, N=30 exp/94 ctrl, Fig. 3D). We also examined the sensitivity of animals in media in which Cl^-^ concentrations were varied to manipulate the strength of the electrochemical gradient supporting anion efflux. We found that increasing this electrochemical gradient by reducing extracellular Cl^-^ (0.8 mM) produced a small increase in the response to stimuli relative to freshwater media (5.2 mM), although this effect was not significant (omnibus p > 0.05, N=30 exp/94 ctrl, Fig. 3E). Conversely, we found that decreasing this gradient by increasing extracellular Cl^-^ (16 mM) reduced the response (27% decrease with 50 μm stimulus, omnibus p < 0.01, N=30 exp/94 ctrl, Fig. 3E) relative to freshwater media. This suppression, but not termination, of lateral line sensitivity is consistent with the reversal potential still favoring anion efflux at this extracellular Cl^-^ concentration (Fig. 1E). Higher Cl^-^ concentrations were examined and similar results were observed (Extended Data Fig. 3), although such salines impose a stronger osmostic stress that may complicate interpretation. Cumulatively, these results provide further support for the role of anion efflux in lateral line mechanotransduction.

### A new hypothesis for lateral line mechanotransduction

The MET channels in hair cells are permeable to both monovalent (K^+^, Na^+^) and divalent (Ca^2+^) cations^7,8^. It has long been speculated that mechanotransduction in lateral line hair cells was mediated by cation influx through the apical membrane, similar to the inner ear. Here we present a new hypothesis: deflection of the hair cell bundle opens MET channels allowing influx of trace amounts of Ca^2+^, which interacts with Ca^2+^-activated Cl^-^ channels, leading to Cl^-^ efflux through the apical membrane that induces membrane depolarization (Fig. 4). Several lines of evidence support this hypothesis, including 1) the electrochemical gradient across the apical membrane is unable to support cation influx induced depolarization, 2) hair cells only require micromolar extracellular Ca^2+^ to function, 3) the hair cells express Ca^2+^-activated Cl^-^ channels, 4) activating these channels induces hair cell depolarization, and 5) blocking these channels reduces the response of animals to mechanosensory stimuli.

**Fig. 4.**
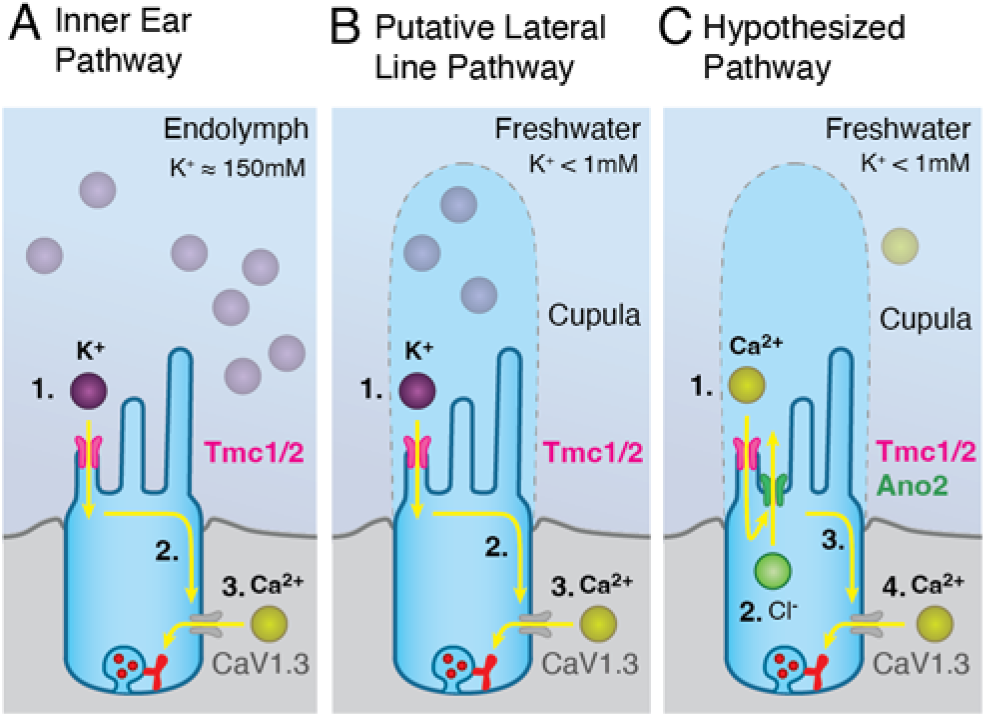
Calcium-activated chloride channels amplify hair cell signaling in environments of low ionic strength. **(A)** *Established inner ear pathway*: Inner ear hair cells are bathed in a K^+^ rich endolymph. 1. As the stereocilia deflect, mechano-electrical transduction (MET) channels open, allowing K^+^ influx into the hair cells. 2. Cation influx initiates membrane depolarization. 3. Voltage-gated calcium channels allow Ca^2+^ influx at the basolateral membrane and subsequent vesicle fusion. **(B)** *Putative Lateral Line pathway*: Similar to (A), except the cupula encapsulating the lateral line hair cells maintains an ionic microenvironment that supports K^+^ influx. (**C**) *Hypothesized Pathway*: The apical membrane of the lateral line hair cells in zebrafish is exposed to external freshwater environments with insufficient cations to directly drive depolarization. We hypothesize that depolarization is mediated by anion efflux through the following processes: 1. Trace amounts of Ca^2+^ influx through MET channels. 2. Calcium-activated chloride channels open. 3. Cl^-^ efflux initiates membrane depolarization. 4. Voltage-gated calcium channels support vesicle fusion.

## Discussion

Anion efflux mediated sensory transduction provides several potential advantages for cells exposed to ion-poor environments^28^. Anion efflux is expected to be robust to fluctuating external conditions, since the intracellular environment provides a well-regulated source of anions^51^ and there is a strong electrochemical gradient supporting efflux across a wide range of external conditions. This intrinsic robustness could allow animals to maintain sensitivity in dynamic environments, without the need for secondary pathways for modulation or auxiliary structures that maintain a stable extracellular microenvironment. Although transduction mechanisms leveraging anion efflux have received much less attention than pathways utilizing cation influx, signal amplification through anion efflux is also found in vertebrate olfactory receptor neurons^22–26^. There, odorant receptors act via a G-protein coupled cascade to increase cAMP, which induces opening of a cyclic-nucleotide-gated cation channel. This leads to an influx of Ca^2+^ and subsequent opening of Ca^2+^-activated Cl^-^ channels (Ano2). This striking similarity between mechanoreceptive and olfactory systems, two seemingly disparate sensory modalities, may suggest convergent evolution in systems exposed to dynamic extracorporeal environments.

The lateral line appears to have evolved in early vertebrates^12^, but it is not clear if these early vertebrates occupied marine or freshwater environments, or both at different life stages^52,53^. In a marine environment, high Na^+^ and Cl^-^ concentrations at the apical surface would easily support cation influx mediated depolarization and make Cl^-^ conductance hyperpolarizing. Here, we show that in freshwater environments transduction would require either the establishment of an ionic microenvironment that can support cation influx or a mechanism based on anion efflux. Understanding the evolutionary origins of lateral line hair cells and how these systems evolved as fishes entered new environments would provide important insights into hair cell physiology and the evolution of sensory systems.

The lateral line system continues to serve as a powerful model for dissecting the basic principles of hearing and balance. However, prior studies by our lab and others have typically examined lateral line physiology in ion-rich extracellular saline that mimics vertebrate blood rather than the ion-poor saline these animals naturally inhabit^54,55^. The use of cation-rich saline would be expected to introduce an artificial cation influx while masking naturally occurring anion efflux. As such, this work highlights the necessity of studying sensory systems in the context in which they evolved in order to decipher their true properties and capacities.

## Methods

### Animals

Experiments were performed on 4-7 day post fertilization (dpf) zebrafish larvae (*Danio rerio*), which were raised in bicarbonate-buffered E3 media (5 mM NaCl, 0.17 mM KCl, 0.33 mM CaCl_2_, 0.33 mM MgSO_4_, 0.35 NaHCO_3_, 0.27 μM methylene blue, pH 7.2-7.4; modified from ^1^) under standard conditions (28°C, 14:10 light:dark) and provided micropowder feed starting at 5 dpf (Microgemma 75, Skretting). All protocols were approved by the University of Florida Institutional Animal Care and Use Committee.

### Cupula ionic composition experiments

Larvae were immobilized by immersion in a paralytic that specifically blocks nicotinic acetylcholine receptors present at the neuromuscular junction (20μL of 1 mg/mL α-bungarotoxin [Sigma Aldrich] in HEPES-buffered E3 [5 mM NaCl, 0.17 mM KCl, 0.33 mM CaCl_2_, 0.33 mM MgSO_4_, 1 mM HEPES, titrated to pH 7.2-7.4])^2^. Larvae were transferred to a coverslip-bottomed recording dish at room temperature (20-22°C), pinned onto a small silicone elastomer pad (5 mm x 1 mm x 1 mm, Sylgard 184) using etched tungsten pins (<50 μm diameter) through the notochord, and then this pad was turned over and pinned to a larger block of silicone elastomer such that the lateral side of the animal faced downward. This lateral-side down configuration enabled imaging of the cupula cross-section for predominantly trunk neuromasts on an inverted confocal microscope. Blood flow was taken as a measure of animal health and only trials with robust blood flow at the beginning and end of the experiment were analyzed. The K^+^ concentration within the cupula was measured by perfusing on media containing the potassium-selective fluorescent indicator (10 μM IPG-4 in HEPES-buffered E3; ION Biosciences), and recording the fluorescence in the cupula and the surrounding media near the stereocilia (Leica TCS SP5 II, HCX PL APO CS 63x/1.2 water objective, 1 frame/sec, excitation: 514 nm, emission: 530 - 574 nm). Measurements were also made 5 μm and 10 μm above the stereocilia and yielded similar results. Fluorescence values were averaged over a manually selected region-of-interest in ImageJ (v1.48; U. S. National Institutes of Health).

The recorded indicator fluorescence inside of the cupula was not substantially different from the low indicator fluorescence present in the surrounding media, suggesting that the K^+^ concentration in the cupula was similar to that of the surrounding media. To ensure that this was not a result of low indicator penetration into the cupula or other issues with fluorescence detection, the relationship between K^+^ concentration and indicator fluorescence was then measured *in situ*. The K^+^ concentration of the media was incrementally increased by washing in media with increased KCl (0.2-60 mM total conc.), and changes in the fluorescence within the surrounding media and cupula were recorded. Again, indicator fluorescence inside of the cupula increased in parallel with the surrounding media, suggesting that both the indicator and K^+^ penetrate into the cupula. To precisely determine the absolute concentration of K^+^ within the cupula under normal conditions (E3 media), these curves were then analyzed using chemical kinetics models. Specifically, fluorescence within the external media was then fit to the Hill equation,

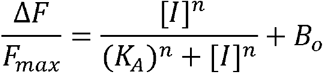

where ΔF is the fluorescence of the indicator minus the background fluorescence without the indicator, F_max_ is the saturated fluorescence of the indicator minus the background fluorescence, B_o_ represents the baseline fluorescence of the indicator, K_A_ is the indicator binding affinity, n is the indicator Hill coefficient, [I] is the K^+^ concentration of the media. Fitting was performed using non-linear least squares optimization (Mathworks, MATLAB 2019b). The computed binding constants for IPG-4 were consistent with previously reported values (K_A_= 5.7 ± 1.5 mM, mean ± S.E.M.^3^). The K^+^ concentration within the cupular matrix under normal conditions (E3 media) was then calculated as

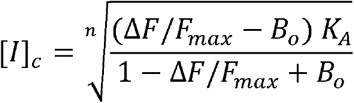

where ΔF/F_max_ is the fluorescence within the cupular matrix as measured in E3 media, normalized by the fluorescence in the cupular matrix at indicator saturation. It should be noted that in this approach the fluorescence values with elevated K^+^ media are used only to calibrate the concentration dependence of the indicator molecule (K_A_, B_o_) and compute the amount of indicator within the cupula (represented by F_max_). Any potential effects of elevated K^+^ on the physiology of the neuromast would not be expected to modify these results, provided that these effects do not alter the chemical partitioning of the ion indicator between the media and cupular matrix. The difference between the K+ concentration in the cupula and the surrounding media was then compared to zero using a two-tailed one-sample t-test.

The Na^+^ concentration was measured using a similar protocol with a different probe (10 μM ING-2 [Ion Biosciences] in HEPES-buffered E3, 5-95 NaCl for calibration, K_A_=21.4 ± 3.1 mM^4^, excitation: 488 nm, emission: 500-550 nm). To record the Ca^2+^ concentration, animals were first imaged in media containing a Ca^2+^ selective indicator (10 μM Fluo5N [Invitrogen] in HEPES-buffered E3, Ka = 13.4 ± 2.3 μM^5^, excitation: 488 nm, emission: 500-550 nm), then the media was replaced by saline solutions with varied Ca^2+^ concentrations (0-2 mM, prepared by varying CaCl_2_, random order). The difference between the Ca^2+^ concentration in the cupula and media was computed for 20 μM Ca^2+^ media rather than E3 media, since this provides a lower baseline and would be more sensitive to deviations. Similarly, to record the H^+^ concentration, animals were imaged in neutral-pH media with a selective probe (10 μM BCECF [Cayman Chemicals] in HEPES-buffered E3 [pH = 7.2], excitation: 496 nm, emission: 477-545 nm), then the media was replaced by saline solutions with varied pH values (first increased to pH=8.4, then decreased to 4.0, prepared by titrating with NaOH or HCl, 5mM MES added to media for solutions with pH < 6.8). Since BCECF is quenched, this calibration curve was fit to the Stern-Volmer equation instead of the Hill equation and F_max_ was taken as the fluorescence at the highest examined pH value, but calculations were otherwise similar. The Cl^-^ concentration within the cupular matrix was measured using the fluorescent indicator MQAE (1 mM in HEPES-buffered E3). In order to provide excitation at the short UV wavelengths required by MQAE, imaging was performed using two-photon microscopy (custom-built system, Zeiss 20x/1.0 objective, Coherent Chameleon Vision II laser, 750 nm excitation, Chroma HQ480/40M emission filter). Since the MQAE could be visualized quickly penetrating through the cupula and to limit the potential for UV-induced damage, the indicator calibration was performed in a separate series of experiments without the animal (0-50mM KCl) rather than with the animal as above. To produce a calibration curve, the fluorescence at each Cl^-^ concentration was normalized by the fluorescence recorded for saline with a Cl^-^ concentration equal to E3 media. The fluorescence measured in the cupula was then normalized by the fluorescence in the surrounding media, and this value was compared to the calibration curve to calculate the concentration of Cl^-^ in the cupula.

### Membrane reversal potentials

The membrane reversal potentials for the MET channel of the lateral line hair cells in different extracellular salines were estimated using a modified constant field equation that includes the contributions of Ca^2+^ (modified GHK equation^6,7^). The examined extracellular concentrations were centered around the values for E3 media, and the intracellular ionic composition was assumed to be similar to other neurons ([Na^+^]_out_ = 5 mM, [Na^+^]_in_ = 5 mM, [K^+^]_out_ = 0.17 mM, [K^+^]_in_ = 130 mM, [Ca^2+^]_out_ = 0.33 mM, [Ca^2+^]_in_ = 100 nM, 28°C, ^8^). The relative permeabilities of MET channels were estimated based on measurements from hair cells in other species: P(Ca^2+^) = 5, P(K+) = 1.15, P(Na^+^) = 1,^9–11^. For comparison, the reversal potential for selective chloride conductance was also estimated using the Nernst equation ([Cl^-^]_in_ = 10 mM, 28°C,^12^). The calculated reversal potentials were compared against the opening potential for the voltage-gated Ca^2+^ channels on the basolateral membrane (CaV1.3, estimated at −50mV,^13–16^).

### Cupula diffusion experiments

Animals were prepared as in the cupula ionic composition experiments. We visualized the precise boundary of the cupula by immersing the animals in fluorescent polystyrene microspheres and allowing them to coat the cupula (200 nm, suncoast yellow, FSSY002, Bang Labs; 1:100 in HEPES-buffered E3, ^17^). The permeability of the cupular matrix was measured by recording the penetration of a negatively-charged fluorophore (6-carboxyfluorescein; Sigma-Aldrich) into the cupula. The fluorophore (10 μM in HEPES-buffered E3) was loaded into a glass pipette (tip diameter: ~5 μm; Model P-97 Flaming/Brown Micropipette Puller, Sutter Instrument Co.) aimed ~20 μm away at the cupula. The pipette pressure was then rapidly increased by opening a valve attached to a pressure source (100 torr, Fluke Biomedical Instruments DPM1B), dye surrounded the cupula, and the change in fluorescence over time was recorded at the stereocilia level (Leica TCS SP5, HCX PL APO CS 63x/1.2 water objective, 20 frames/sec, excitation: 488 nm; emission: 500 - 550 nm). Values for ΔF/F_max_ were computed by subtracting the fluorescence prior to dye release, and then normalizing by the maximum fluorescence following release. The time constant for the increase in dye fluorescence was fit to an exponential waveform

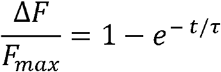

where F_max_ is the maximum fluorescence, τ is the time constant, and t is time. Fitting was performed using non-linear least-squares fitting (Mathworks, MATLAB R2020b). Relative time constants were calculated as the ratio of the time constant in the cupula divided by the time constant in the surrounding media for the same trial. Differences were identified by comparing this relative time constant to one using a two-tailed one-sample t-test.

### Computational cupula diffusion model

Simulations of the fluorophore diffusion experiments were performed by modeling the cupula as a cylinder (13 μm diameter, 40 μm height, ^17^) of uniform material with an isotropic diffusion coefficient. Since the pipette jet rapidly replaced the fluorophore solution at the periphery of the cupular during experiments, the fluorophore concentration was assumed to be uniform over the exposed surface of the cupula and increase following an exponential waveform with parameters estimated from the imaging data, see cupula diffusion experiments). Diffusion is a linear process and the absolute fluorophore concentration does not alter the dynamics, so the concentration was arbitrarily selected to ramp from zero to one. The cylinder was discretized using an all-tetrahedral mesh and Fick’s equation was solved for this geometry using a commercial finite element method solver (Comsol 6.0). A sensitivity analysis was performed to ensure that mesh density and time stepping were adequate. Although the central problem is axisymmetric, a 3-dimensional mesh was used as some sensitivity analyses included non-axisymmetric convective terms. This model was then solved for a logarithmic series of cupular matrix diffusion coefficients surrounding the value recorded for the fluorophore in free water (1×10^−12^ to 1×10^−9^ m^2^/s, D_6CF_ = 0.487×10^−9^, ^18^), and the predicted temporal profile of the fluorophore at the center of the cupula was calculated for each condition. The cupular diffusion coefficient was then estimated by comparing the time constants from these simulations to the time constants obtained from the imaging experiments.

### Computational Nernst-Planck model

We performed a series of computational studies to determine how K^+^ secretion at the apical surface of the neuromast would affect K^+^ concentrations within the cupula. The cupula was modeled as a cylinder (13 μm diameter, 40 μm height, ^17^) immersed in an infinitely large electrolyte bath (spherical bubble with radius of 100 μm explicitly modeled, surrounding volume modeled using infinite element domain). The ion currents and concentrations within the cupula and surrounding media were modeled using the full Nernst-Planck equations subject to the electroneutrality approximation and solved using a commercial finite element method multiphysics package (Comsol 6.0). In addition to water, the electrolyte included K^+^, Na^+^, and Cl^-^ ions and the self-diffusion coefficients for each ion in the surrounding media were taken from previously reported values (K^+^: 1.89×10^−9^, Na^+^: 1.33×10^−9^, Cl^-^: 2.06×10^−9^ m^2^/s, ^19,20^). The model was solved for two configurations: the most probable scenario in which the ion diffusion coefficients in the cupular matrix are equivalent to those in water, and a conservative model in which the diffusion coefficient of each ion was scaled down to match the lower bound of the 95% CI for the relative diffusion coefficient obtained in the 6-CF experiments (6.9% of free water). The electrical mobility of each species was estimated using the Nernst-Einstein relation. The model was solved using a steady-state axisymmetric system, the cupula and explicit media domain were meshed using free triangular elements, the infinite element domain was meshed with mapped quadrilateral elements, and a sensitivity analysis was used to confirm that the mesh density was sufficient. The boundary conditions were set such that at infinity the electrical potential was zero and the ion concentrations were equal to those of E3 media (0.17 mM K^+^, 5 mM Na^+^, 5.17 mM Cl^-^), the surface of the body was impermeable to all ions, and the base of the cupula secreted K^+^ ions at a fixed rate. Simulations were performed for a logarithmic series of K^+^ secretion currents spanning a wide range (1×10^−10^ - 1×10^−6^ A), and K^+^ concentrations at the base of the cupula and in the surrounding media were computed for each current value. Results were qualitatively similar for the water-like diffusion configuration and the conservative configuration with scaled down diffusion coefficients, although the conservative configuration yielded higher K^+^ concentrations within the cupula as expected (reaching a K^+^ concentration of 10.2 mM at the apical surface requires 56 nA for a diffusion coefficient equal to water, and 4.8 nA for a diffusion constant at the lower bound of the 95% confidence interval).

### Calcium imaging

We examined the responses of the lateral line hair cells to mechanical stimuli in environmental salines with varying ionic compositions. Larvae (5 −7dpf) expressed the membrane-localized fluorescent calcium indicator gCaMP6s-CAAX in hair cells (*myo6b:gCaMP6s-CAAX*, ^21^), and were immobilized and imaged as described above (in HEPES-buffered E3). A glass bead (5 μm) attached to a piezoelectric transducer (30V300, Piezosystem Jena) was positioned approximately 10 μm anterior from the distal end of the neuromast kinocilia^22,23^. During each trial, a stimulus program (Clampex 10.1) controlled via a low-noise digitizer (Digidata 1440A) elicited three sweeps of sinusoidal movement (5 Hz) for 20 seconds that were preceded and followed by inactivity of equal duration. The glass bead vibrated, deflecting the cupula in the anterior-to-posterior direction. Cupula deflection was monitored by measuring the displacement distance of the kinocilia tips (~10 μm). Mechanosensitive Ca^2+^ responses were measured within the hair cells (Leica TCS SP5 II, HCX PL APO CS 63x/1.2 water objective, 5 frames/sec, excitation: 488 nm, emission: 501 - 583 nm). Between trials, the perfusate bath was replaced using media with decreased total Ca^2+^ concentration (20 μM Ca^2+^: HEPES-buffered E3 with reduced CaCl_2_, or 0 μM Ca^2+^: HEPES-buffered E3 with CaCl_2_ replaced by equivalent concentration of MgSO_4_). Preliminary experiments performed in media containing EGTA (0.2 mM) yielded similar results to 0 μM Ca^2+^ trials, suggesting these solutions contained negligible residual calcium. Responses to stimuli were quantified by finding the mean fluorescence within manually selected regions of interest (ROI) for individual hair cells (ImageJ). Fluorescence values were then used to compute ΔF/F by subtracting and then normalizing by the baseline fluorescence computed from the mean fluorescence within the cell prior to stimulation. Responses were detected by comparing the time-average of the ΔF/F during stimulus periods to non-stimulus periods using a paired two-tailed t-test. Changes in response amplitude were identified by comparing the time-average during stimulus periods between treatment groups (20 μM Ca^2+^ and 0 μM Ca^2+^ media) and the control (E3 media) using a paired two-tailed t-test.

A similar protocol was used to examine the responses of hair cells to a calcium-activated chloride channel agonist. Baseline activity was recorded for 30 s in freshwater media (E3 saline), then perfused with Eact (50 μM in 0.1% DMSO final conc., ^24^). We monitored changes in fluorescence before, during, and after perfusion of Eact (0.8 frames/sec, excitation: 488 nm, emission: 500 nm - 600 nm). Relative changes in fluorescence were quantified using regions of interest around responsive hair cells. Mean ΔF/F during the 30 sec prior to Eact exposure was then compared to the mean response for 60 sec following Eact exposure using a paired two-tailed t-test.

### Behavioral assays

The behavioral response of zebrafish larvae (5-7 dpf, AB strain) to mechanical stimuli was assessed using a custom-developed behavioral assay. This assay was conducted in an enclosure that maintains a constant internal temperature (28°C, using Fisher Scientific IsoTemp 6200 R28) and isolates experiments from ambient light and noise. Animals were transferred into a dish containing the experimental media (38 mm diameter, 6 mm height) to incubate (20 min for all trials except neomycin, which incubated for 1 h), then transferred into the behavioral chamber and allowed to acclimate briefly (5 min) prior to experiments. The behavioral chamber was designed to operate as an oscillatory Couette cell, which produces shear stress in the fluid but not pressure waves ^25^. This chamber was laser cut from acrylic sheet and provided a cylindrical raceway (35 mm outer diameter, 25 mm inner diameter, filled to depth of 5 mm) that was suspended by flexible struts. These flexible struts functioned as a flexure bearing system, allowing oscillatory rotations of the raceway about the cylindrical axis while preventing translation or other rotations. Oscillatory rotations were driven using a fourth, rigid strut that was affixed with adhesive to the center dome of a speaker (Soberton SP-3114Y), which was connected to an audio amplifier (Adafruit 987, MAX98306). The chamber was diffusely illuminated from above with white light (ST-WP-5050-DL-RL, TheLEDLight.com) to maintain normal swimming behavior.

Stimulus generation was controlled via a custom-written MATLAB script (Mathworks, R2020b) and a data acquisition system (National Instruments, PCIe-6323). Each experiment consisted of a baseline activity recording period (300 s) followed by a series of 35 trials with stimuli. Each trial began with a 2 s mechanical stimulus (20 Hz, 5 amplitudes with 7 replicates each, randomly ordered) followed by a randomized interstimulus interval (mean: 120 s, std. dev.: 20 s, min: 60 s, max: 180 s). The magnitude of the stimulus was calibrated by imaging the rigid strut of the raceway using a high-speed camera system (Phantom Miro 340, 200 fps, Samyang 1.4/85mm lens, 68 mm of extension tubes) and computing the displacement with sub-pixel precision using a custom-written MATLAB script. Reported stimulus amplitudes represent the displacement amplitude at the outer wall of the raceway.

The movement of the animals was continuously recorded and used to assess the response to the mechanical stimulus. A sheet of rear screen projection material (GooScreen BlackMax 1950) was positioned beneath the behavioral arena and illuminated with near-infrared light (850nm, ThorLabs 850L3). Animals were then imaged using a NIR-sensitive camera positioned above the arena (30 frames/sec, Point Grey GS3-U3-41C6NIR) equipped with a NIR-transmissive lens (Schneider 50mm Xenoplan, 1001976) and visible light blocking filter (Lee Filter #87). During each stimulus period, a small NIR LED positioned within the field of view was automatically illuminated, and this light was used to synchronize the stimuli with the recorded video to within one frame. The position of the larvae in each image frame was calculated using previously developed MATLAB scripts (https://bitbucket.org/jastrother/larval_proving_grounds).

The response to each stimulus was scored based on the induced change in the swimming speed of the larvae. In order to quantify each response in a way that was independent of the average swimming speed of the animal, each trial was scored as a response if the stimulus elicited a change in the swimming speed that fell outside of the 95% confidence interval computed for similar stimulus-free periods. Specifically, the average swimming speed of the animals was computed for 3 time windows: a first reference window (500 ms duration, ending 10 s prior to the stimulus), a second reference window (500 ms duration, ending 5 s prior to the stimulus), and a response window (500 ms duration, starting with the stimulus). The difference between the first and second reference window was aggregated and used to compute the cumulative probability distribution of the spontaneous swimming speed change during a stimulus-free period for each individual. The difference between the second reference window and the response window was compared to this distribution, and each trial was marked as a response if it fell below the 2.5% percentile or above the 97.5% percentile. Since this approach yields an expected false positive rate of 5%, the calculated response probabilities were offset by the same amount.

Differences in the response probability between experimental and control groups were detected with two different statistical analyses. First, multinomial probit regression was used to implement an omnibus test that detects differences between the experimental and control groups while integrating information across stimulus intensities (Mathworks, MATLAB R2020b). Second, pairwise differences between the experimental and control groups at specific stimulus intensities were detected using Fisher’s exact test with false discovery rate control implemented using the Benjamini-Hochberg procedure (Mathworks, MATLAB R2020b). Plots include error bars representing S.E.M. values, where the variance was calculated using the properties of binomial distribution [p (1-p)].

The following conditions were examined: “HEPES-buffered E3 media” (5 mM NaCl, 0.17 mM KCl, 0.33 mM CaCl_2_, 0.33 mM MgSO_4_, 1 mM HEPES, titrated to pH 7.2-7.4 with HCl/NaOH), “200 μM Neomycin media” (200 μM neomycin sulfate in HEPES-buffered E3 media), “50 μM niflumic acid” (50 μM niflumic acid in HEPES-buffered E3 media with 0.05% DMSO), “3x Ionic Strength media” (NaCl and KCl increased 3x relative to E3: 15 mM NaCl, 0.51 mM KCl, 0.33 mM CaCl_2_, 0.33 mM MgSO_4_, 1 mM HEPES, titrated to pH 7.2-7.4 with HCl/NaOH), “20x Ionic Strength media” (NaCl and KCl increased 20x relative to E3: 100 mM NaCl, 3.4 mM KCl, 0.33 mM CaCl_2_, 0.33 mM MgSO_4_, 1mM HEPES, titrated to pH 7.2-7.4 with HCl/NaOH), “0.8 mM Cl^-^ media” (replace NaCl with Na-Gluconate in E3: 5 mM Na-Gluconate, 0.17 mM KCl, 0.33 mM CaCl_2_, 0.33 MgSO_4_, 1 mM HEPES, titrate to pH 7.2-7.4 with citric acid/NaOH), “16 mM Cl^-^ media” (major cations matched to E3, approximately isosmotic to 3x Ionic Strength media: 5mM NaCl, 10.3 mM NMDG-Cl, 0.17 mM KCl, 0.33 mM CaCl_2_, 0.33 MgSO_4_, 1mM HEPES, pH 7.2-7.4), “104 mM Cl^-^ media” (major cations matched to E3, approximately isosmotic to 20x Ionic Strength media: 5 mM NaCl, 98.22 mM NMDG-Cl, 0.17 mM KCl, 0.33 mM CaCl_2_, 0.33 mM MgSO_4_, 1mM HEPES, titrated to pH 7.2-7.4 with HCl/NaOH), and “20 μM Ca^2+^ media” (replace Ca^2+^ with Mg^2+^ in E3: 5 mM NaCl, 0.17 mM KCl, 20 μM CaCl_2_, 0.31 mM MgCl_2_, 0.33 mM MgSO_4_, 1 mM HEPES, titrated to pH 7.2-7.4 with HCl/NaOH). The HEPES-buffered E3 media served as the control for all experimental groups, and control trials were run in parallel with experimental trials. Since the responses from control trials exhibited very little inter-clutch variation, data from control trials was pooled when performing statistical analyses. At least 3 clutches of embryos were used for all conditions, and parents were randomly selected from a genetically diverse population.

### Analysis of transcriptomic data

In order to identify gene products that may be contributing to the mechanotransduction process, we conducted a survey of previously published lateral line neuromast scRNA-seq data ^26^. A MATLAB script (Mathworks, R2020B) was written that performed the following steps: cells that express the hair cell marker *tmc2b* were selected, the transcripts expressed by each selected cell were filtered to include genes annotated with a relevant list of ontology terms (“ion transport”, “ion channel activity”, “gpcr activity”, “chemical synaptic transmission”, “hormone activity”, “neuropeptide hormone” for *Danio rerio* in AmiGo2^27^ and a manually curated listed of similar genes not captured in the gene ontology database (based on searches of zfin.org^28^), and the resulting list was then reviewed to identify genes with known anion efflux activity. This analysis was performed exclusively for hypothesis generation. To avoid issues with pseudoreplication, further examination of the genes of interest was performed in independent experiments (e.g., *in situ* hybridization) rather than a statistical analysis of the same transcriptome dataset.

### *In situ* hybridization

The expression of *ano2b* was examined using HCR RNA-FISH^29^ following protocol based on the manufacturer’s directions. Briefly, larvae (AB strain, 4 dpf) were cold anesthetized and fixed with 4% PFA in PBS (overnight at 4°C), washed with PBS (3x, 5 min, room temperature [RT]), dehydrated with a PBS/MeOH series (75/25%, 50/50%, 25/75%, 0/100%, 5 min each at RT), frozen (1 hr at −20°C), rehydrated with a MeOH/PBST series (75/25%, 50/50%, 25/75%, 0/100%, 5 min each at RT, PBST: PBS with 0.1% Tween 20), incubated in hybridization buffer (20 min at 37°C), hybridized with *ano2b* probes (20 nM, overnight at 37°C), washed with probe wash buffer (4x, 15 min, 37°C), washed with 5X SSCT (2x, 5 min, RT; SSCT: SSC with 0.1% Tween 20), incubated in amplification buffer (30 min, RT), incubated with snap-cooled amplifiers in amplification buffer (60 nM each, 18 h, RT), washed with 5X SSCT (2x 5 min, 2x 30 min, 1x 5 min, RT), labeled with DAPI (30 min, 1 μM in 5X SSCT, RT), and imaged on a confocal microscope (Leica TCS SP5, HCX PL APO CS 63x/1.2 water objective for hair cells, HC PL Fluotar 20x/0.5 objective for others, DAPI: excitation = 405 nm, emission = 415 - 485 nm, AlexaFluor 647: excitation = 633 nm, emission = 695 - 765 nm). Images were collected of superficial neuromasts, olfactory epithelium, and the CNS. Images were also collected from randomly selected regions along the trunk adjacent to superficial neuromasts, in order to verify that the observed labeling of superficial neuromasts was not a product of non-specific surface labeling.

## Acknowledgments

We thank K. Kindt for providing plasmids and transgenic zebrafish embryos, J. Ryan for his insights on transcriptome analysis, and the UF Center for Taste and Smell for its support. Funding provided by: National Science Foundation (IOS1932707 to JAS; IOS1257150, IOS1855956, & IOS1856237 to JCL), Paul G. Allen Frontiers Group (to JAS), and National Institutes of Health (R01DC010809 to JCL).

## Author contributions

ETL, YVB, JCL, and JAS conceived the study and designed the methodology. ETL, YVB, and JAS conducted imaging experiments and analyzed the data. BCR and JAS conducted behavioral experiments and analyzed the data. ETL, YVB, JCL, and JAS interpreted the results. ETL, BCR, JCL, and JAS contributed to the first draft of the manuscript, and all authors contributed to reviewing and editing the manuscript.

## Additional Information

Correspondence and requests for materials should be addressed to James Strother.

## Data and materials availability

All data and code used for analysis will be made available upon request.

## Extended Data Figures

**Extended Data Fig. 1:**
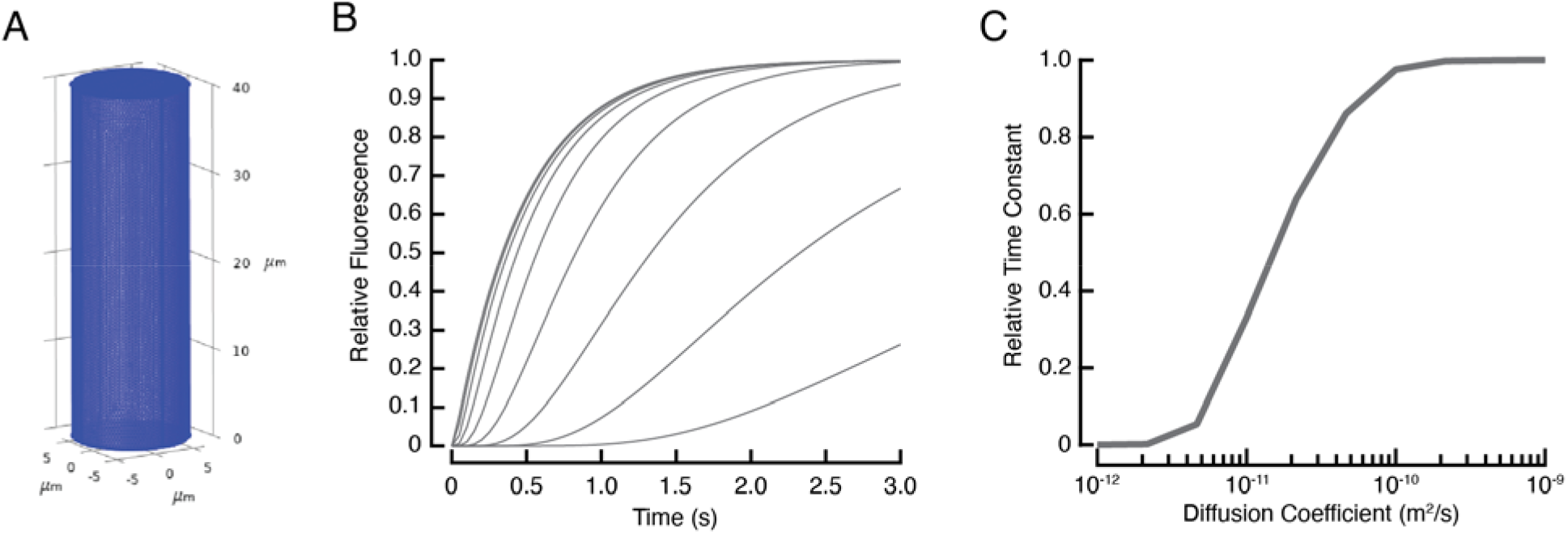
Computational simulations support cupula diffusion experiments. **(A)** Mesh used for finite element modeling. **(B)** Simulated time series for a position just above the base of the cupula (8 μm) for a range of diffusion coefficients (1×10^−12^, 2.15×10^−12^, 4.64×10^−12^, 1×10^−11^, 2.15×10^−11^, 4.64×10^−11^, 1×10^−10^, 2.15×10^−10^, 4.64×10^−10^, 1×10^−9^ m^2^/s). **(C)** Simulated relative time constant as a function of the modeled diffusion coefficient. Comparison of experimental and modeling results indicate that the diffusion of the charged fluorophore 6-CF through the cupular matrix is similar to that through water (D = 0.034×10^−9^ - 0.487×10^−9^ m^2^/s, 6.9%-100% of water, 95% CI), with the lower bound of the estimate limited by the achievable dye injection speed of the experimental apparatus.

**Extended Data Fig. 2:**
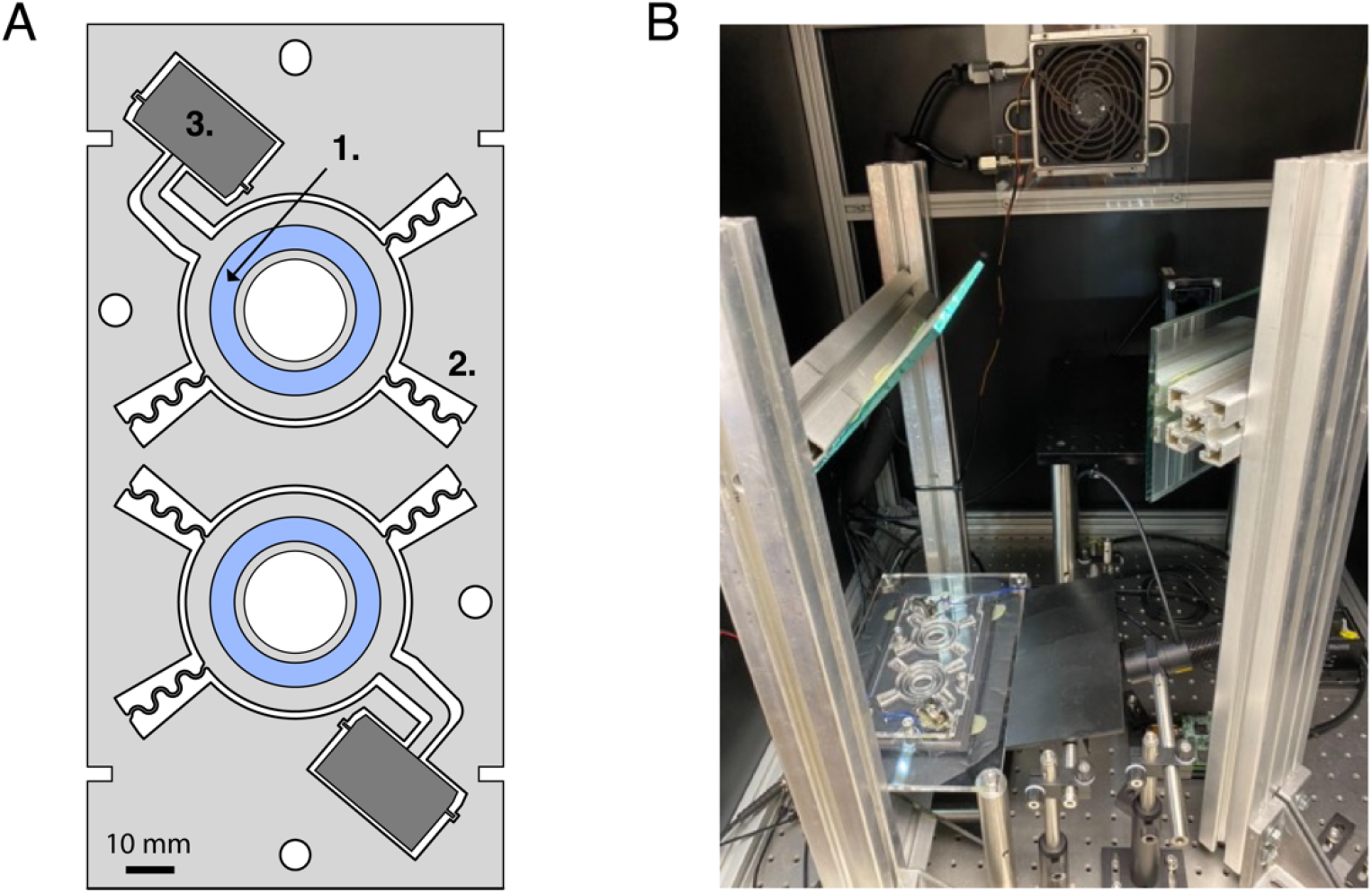
Apparatus used for behavioral assays. **(A)** Outline of the behavioral arena, showing circular racetrack in which larva swims [1], which is suspended by flexure bearings [2] and driven by a speaker [3]. Each apparatus contains two chambers that are operated in parallel. **(B)** Image of behavioral assay in temperature controlled enclosure.

**Extended Data Fig. 3:**
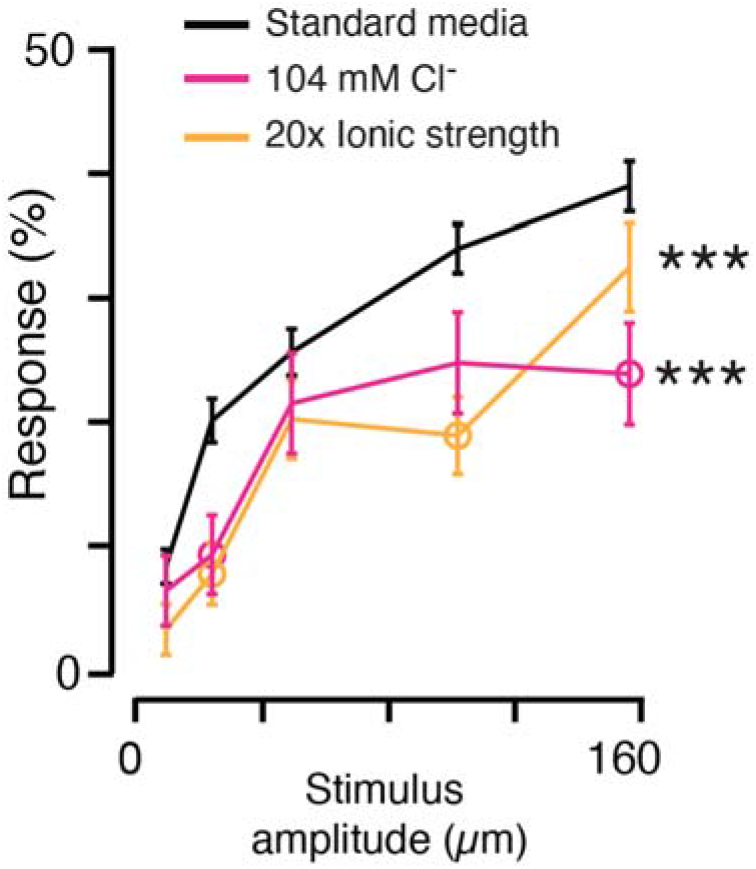
Zebrafish larvae display reduced lateral line sensitivity in environmental saline solutions with higher ionic strength. The response probability of zebrafish larvae to mechanical stimuli of different intensities in various environmental saline solutions, as measured using the oscillatory Couette behavioral assay. Asterisks indicate statistically significant differences integrated across stimulus intensities (*p<0.05, **p<0.01, ***p<0.001), open circles represent significant pairwise comparisons (p<0.05), and results are shown as mean ± SEM.

**Extended Data Fig. 4:**
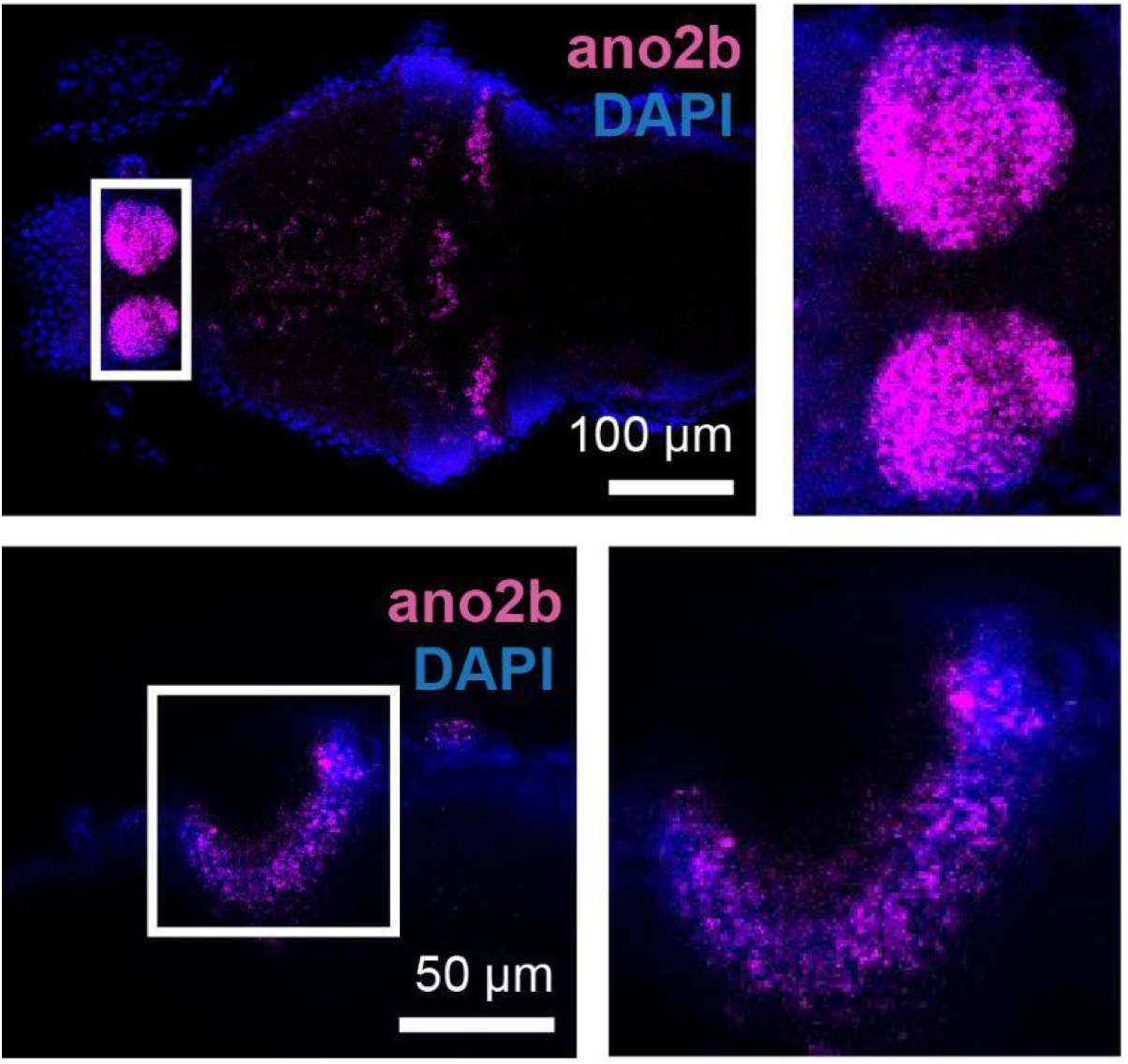
*In situ* hybridization for *ano2* labels the dorsal habenula (top, magnified right) and the olfactory epithelium (bottom, magnified right).

